# Long-range Notch-mediated tissue patterning requires actomyosin contractility

**DOI:** 10.1101/259341

**Authors:** Ginger L Hunter, Li He, Norbert Perrimon, Guillaume Charras, Edward Giniger, Buzz Baum

## Abstract

Dynamic, actin-based protrusions function in cell-cell signaling in a variety of systems. In the dorsal thorax of the developing fly, basal protrusions enable non-neighboring epithelial cells to touch, extending the range over which Notch-mediated lateral inhibition can occur during bristle patterning. Given that actin-based cell protrusions can exert mechanical forces on their environment and Notch receptor activation is mechanically sensitive, how might cytoskeletal contractility contribute to Notch signaling? We identify a pool of basal non-muscle myosin II (myosin II) that regulates protrusion dynamics, promotes Notch signaling, and is required in signal sending and receiving cells for Notch-dependent patterning. We show that interactions between protrusions are extensive and subject to actomyosin contractility. The effects of reducing myosin II activity are more pronounced for protrusion-mediated signaling than for signaling at lateral cell contacts. Together, these results reveal a role for actomyosin contractility in Notch activation, signaling, and patterning in a developmental context.

## Introduction

The ability of cells to signal at a distance through the extension of long, thin, cellular protrusions (e.g., cytonemes) has changed the way that we think about morphogen-based cell-cell signaling in development and disease (Kornberg and Roy, 2014). Across several species and signaling paradigms, it has been observed that signal sending cells use protrusions to directly contact signal receiving cells at a distance, often in other tissues, and elicit a signal response (Bischoff et al., 2013; Cohen et al., 2010; Hamada et al., 2014; Huang and Kornberg, 2015; Sagar et al., 2015). Many types of these protrusions have been shown to carry morphogenetic signaling proteins (e.g., Wnt, Shh, TGFß), and interfering with protrusion lengths can disrupt developmental signaling, leading to changes in pattern formation. One example of a requirement for protrusion mediated signaling is the patterning of mechanosensory bristles on the dorsal thorax of the fruit fly Drosophila melanogaster. At the onset of pattern development in the pupal stages, the dorsal thorax (or notum) tissue is composed of epithelial cells. Over time, these cells undergo a cell fate decision to either acquire a pro-neural, sensory organ precursor (SOP) fate, or to remain epithelial cells. This decision requires lateral inhibition signaling, mediated by the transmembrane ligand, Delta, and its receptor Notch (Artavanis-Tsakonas and Simpson, 1991). In this system, the activation of Notch is induced via contact with another cell expressing Delta; this represses the pro-neural fate so that Notch activated cells tend to remain epithelial in character. Conversely, the maintenance of low levels of Notch activation during this window of development allows the expression of pro-neural genes, causing cells to take on a mechanosensory bristle fate (Furman and Bukharina, 2008). Since Notch-Delta signaling requires direct contact between cells expressing membrane-bound ligand and receptor pairs, for many years it was unclear how this process could give rise to the relatively sparse bristle pattern observed in the wildtype notum (Hartenstein and Posakony, 1990). A possible solution to this conundrum was suggested by studies showing that cells engaging in Notch-Delta lateral inhibition signaling extend long, dynamic protrusions from their basal surfaces. By combining mathematical modeling with live imaging and perturbation experiments, it was possible to show that these dynamic basal protrusions extend the range over which cells can contact one another to pattern the notum (Cohen et al., 2010; De Joussineau et al., 2003; Hadjivasiliou et al., 2016).

The ability of cells to engage in Notch signaling via cellular protrusions is called long-range lateral inhibition. Understanding this type of long-range lateral inhibition requires an understanding of the molecular regulation of protrusion dynamics and mechanics. The protrusions found in notum epithelial cells are formed as the result of actin filament formation downstream of Rac, SCAR and Arp2/3 complex activity (Cohen et al., 2010; Georgiou and Baum, 2010). This is relevant because there are several points of intersection between actin cytoskeletal biology and Notch signaling. First, protrusion dynamics were suggested to be important for Delta-Notch dependent patterning in this system (Cohen et al., 2010). Second, Notch signaling in many systems depends on actin-dependent endocytosis (Mooren et al., 2012; Musse et al., 2012). At the same time, Notch signaling influences the organization of filamentous actin networks (Basu and Proweller, 2016; Le Gall et al., 2008; Major and Irvine, 2005). Finally, recent work has implicated mechanical force in the activation of Notch receptor. Thus, in cell culture, pulling forces of between 3.5 and 5.5 pN are required to expose the S2 site of Notch required for cleavage and Notch activation. This has been proposed to explain the requirement for endocytosis in Notch signaling – since Delta endocytosis is likely to generate sufficient force to induce Notch cleavage and activation (Luca et al., 2017; Meloty-Kapella et al., 2012; Ploscariu et al., 2014; Shergill et al., 2012; Wang and Ha, 2013). However, the actomyosin cytoskeleton that underlies the dynamics of actin-based protrusions can itself generate contractile forces when crosslinked with molecular motors such as non-muscle myosin II (hereafter, Myosin II). Thin actin-based projections are known to exert pulling forces on their environment up to 1 nN (Bornschlogl, 2013; Leijnse et al., 2015). This raises the question of whether the actomyosin cytoskeleton contributes directly to the activation of Notch and to lateral inhibition signaling (Ferguson et al., 2017; Fritzsche et al., 2017; Koster et al., 2016).

Here, we investigate the role of actomyosin contractility on long-range Notch signaling during the patterning of SOP cells in the Drosophila pupal notum using a combination of quantitative live cell imaging and genetic manipulations. By genetically and pharmacologically modulating Myosin II activity in vivo and in a cell culture model of lateral inhibition signaling, we demonstrate the presence of actomyosin-based forces between basal cellular protrusions in an epithelium, and show that a robust Notch response requires Myosin II-mediated contractility in both signal sending and receiving cells. As a result, decreased Myosin II activity is also associated with defects in patterning.

## Results

### Myosin II activity is required for robust Notch signaling

Myosin II is a key contributor to cellular mechanical force production during development via its interactions with filamentous actin in epithelial cell sheets (Curran et al., 2017; Franke et al., 2005; Munjal and Lecuit, 2014). In order to determine whether actomyosin contractility is required for long-range, protrusion-based, lateral inhibition signaling during notum pattern formation, we asked how decreasing actomyosin tension affects the activity of a transcriptional reporter of Notch signaling, NsfGFP (Figure 1A-B) (Hunter et al., 2016). We measured the average accumulation of GFP over time as a reporter of Notch activity (hereafter, rate of Notch response). We used the GAL4/UAS expression system to perturb the function of Myosin II, a hexameric motor protein whose heavy chain is encoded by the Drosophila gene zipper, and regulatory light chain (RLC) is encoded by spaghetti squash (Karess et al., 1991; Kiehart et al., 1989). Previous work showed that loss of function mutations and/or expression of dominant negative derivatives of zipper or RLC leads to phenotypes consistent with decreased cortical tension (Franke et al., 2005; Vasquez et al., 2014). Since animals homozygous mutant for null alleles of zipper (or squash) are not viable to pupariation, we used tissue-specific expression of constructs designed to perturb Myosin II function in specific populations of cells to assess the impact of Myosin II on the morphology and dynamics of basal protrusions in the notum. These included zipper^DN^, a motor-less heavy chain protein that binds and sequesters wildtype heavy chain, thus lowering contractility (Franke et al., 2005), a non-phosphorylatable variant of the RLC, squash^AA^ (Vasquez et al., 2014), or RNAi-mediated silencing of Rho kinase (ROK) an upstream activator of Myosin II contractility (Verdier et al., 2006).

**Figure 1.**
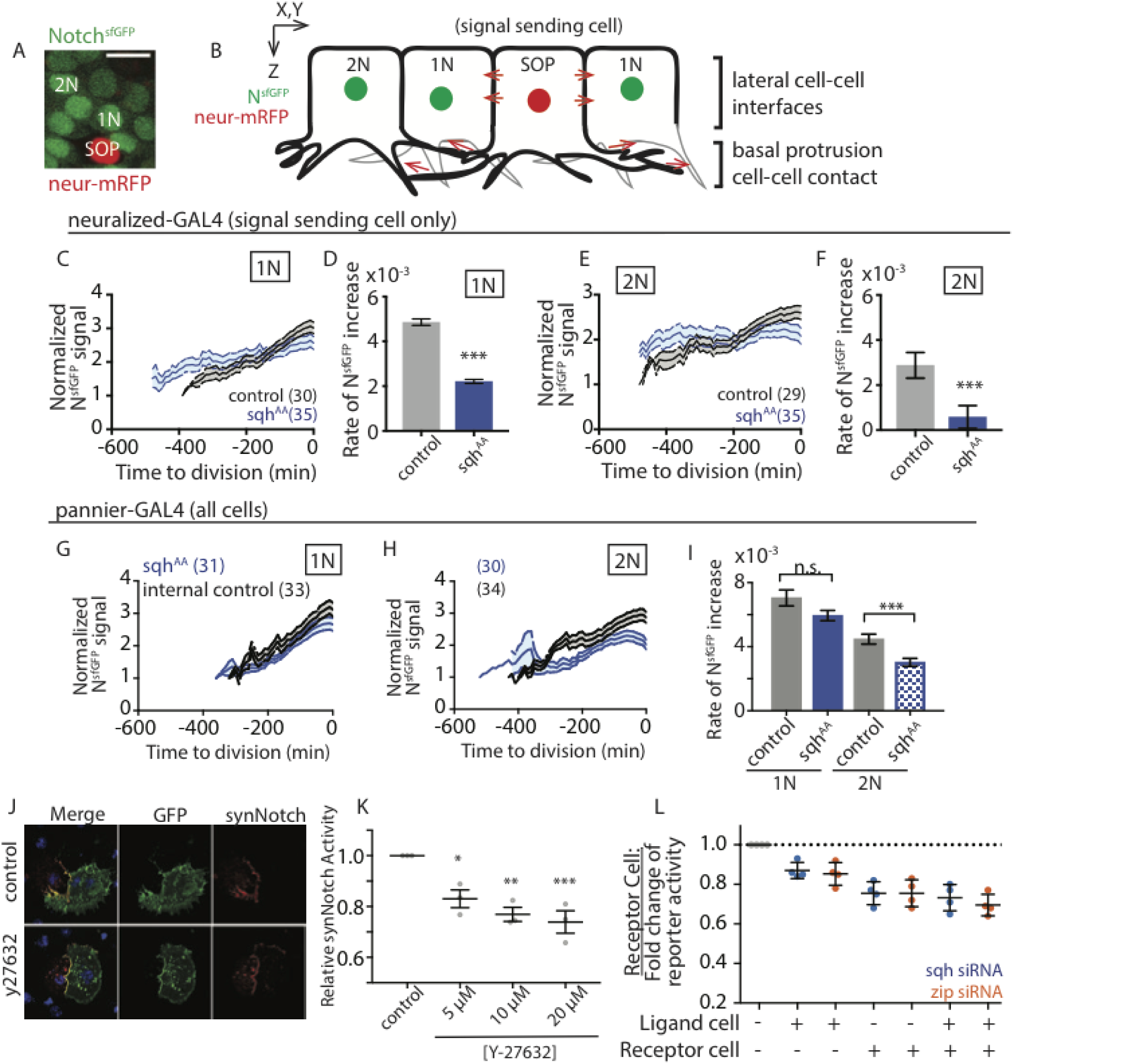
Myosin II activity modulates Notch response in notum epithelial cells. (A) The Notch reporter NsfGFP in 1N, epithelial cell neighbors adjacent to SOP cells; 2N, epithelial cell neighbors at least one cell diameter away from any SOP cell. Neur-mRFP (neuralized- H2BmRFP) is expressed to label SOP cell nucleus, scale bar = 10*µ*m. (B) Cartoon model of adjacent Notch signaling via lateral cell-cell contacts and protrusions vs cells signaling via only basal protrusion contacts. (C-F) Notch response (mean ± SEM) in wildtype epithelial (C) adjacent or (E) distant neighbors to SOP cells expressing UAS-sqh^AA^ (blue) or UAS-LifeActRuby (black) under neur-GAL4 driver. (D, F) Mean ± SEM linear regression slopes for data averaged in (B, D). ***, p ≤ 0.001 by unpaired t-test. (G-I) Notch response in tissue where both SOP and epithelial cells express UAS-sqh^AA^ under the pnr-GAL4 driver. Internal controls were measured outside the pnr domain. Expression of sqh^AA^ does not affect NsfGFP levels (mean ± SEM) in (G) adjacent but does decrease NsfGFP in (H) distant cells compared to internal controls, and only decreases the rate of Notch response in distant cells (I, mean ± SEM). ***, p ≤ 0.001; ns, not significant by unpaired t-test. (n) = total number of nuclei measured, N = 3 nota analyzed per genotype. (J) Images of heterogeneous cell culture to measure synthetic Notch response to Myosin II activity. Cells in green express ligand, those in red express receptor. (K) Relative levels of synNotch activation in response to increasing concentrations of Y-27632. Mean ± SEM, *, p≤0.05, **, p≤0.01, ***, p≤0.001, by ANOVA with multiple comparisons. (L) Relative levels of synNotch activation in response to siRNA against zipper (orange) or squash (blue), in the presence of DMSO. + indicates transfection with siRNA, – indicates transfection with control siRNA targeting the white gene. Mean ± SEM for 4 experimental repeats shown. See also Figure S1.

We expressed UAS- Squash^AA^ or UAS-ROK RNAi in order to reduce Myosin II activity in signal sending cells alone (SOP cells, using neuralized-GAL4) or in both signal sending and receiving cells (using pannier-GAL4)(Amano et al., 1996; Kasza et al., 2014). Strikingly, reducing Myosin II activity in signal sending cells was sufficient to decrease the rate of the Notch response in adjacent, wildtype neighbor cells (Figure 1C-D; S1A-B), and in distant neighbors that are only able to receive signal via protrusion-based interactions (Figure 1B; E-F; S1C). Similar effects were observed when Myosin II activity was compromised in both signal sending and receiving cells simultaneously (Figure 1G-I; S1D-E). In this case, the decrease in Notch response was more profound in distant neighbors, suggesting a greater requirement for Myosin II for lateral inhibition signaling between cells that only contact one another via long, basal protrusions.

To test whether this represents a general role for Myosin II in Notch signaling, we turned to a cell culture model of lateral inhibition (Gordon et al., 2015). In this system, one can mix populations of Drosophila S2R+ cells expressing either a synthetic Notch ligand or receptor in the presence or absence of pharmacological inhibitors of Myosin II activity (Figure 1J; (Ishizaki et al., 2000) or dsRNA mediated knockdown of Zipper or Squash. Once cells form contacts, a luciferase-based transcriptional reporter can be used as a measure of Notch activity. Importantly, while acute treatment of the ROK inhibitor Y-27632 altered S2R+ cell shape, it did not change expression levels of ligand or receptor (Figure 1J; S1F). Nevertheless, ligand-induced Notch signaling in this system was reduced by Y-27632 treatment in a dose-dependent manner (Figure 1K). The role of cortical actomyosin-based tension in Notch signaling was confirmed using dsRNA-mediated knockdown of Zipper or Squash expression (Figure 1L). Moreover, by mixing control and dsRNA treated cells, we were able to show that the maximal Notch response requires Myosin II in both signal sending and receiving cells. These findings support a role for actomyosin contractility in driving robust Notch response.

### Loss of Myosin II activity disrupts bristle patterning

Are these changes in the Notch response sufficient to induce changes in lateral inhibition mediated patterning? To test whether this was the case, we measured the spacing between SOP cells in animals with altered levels of Myosin II (Figure 2A). In this system, defects in Notch-Delta signaling at adjacent contacts are expected to lead to the formation of SOP cell clusters, whereas a failure to signal via protrusions is likely to compromise SOP spacing. The expression of dominant negative Myosin II, a treatment that leads to a strong reduction in Myosin II activity, was associated with decreased spacing between SOP cells in rows, but not with SOP cell clustering (Figure 2A-C). Similar results were seen in flies expressing squash^AA^ in signal sending cells, which led to a reduction in pattern spacing. Furthermore, the expression of squash^AA^ throughout the notum led to a decrease in bristle spacing (Figure 2D-E) – a phenotype that was most evident in bristle row 1 – but again, not to the formation of SOP cell clusters. Together, these data suggest that there is a specific effect of actomyosin-based tension in long range, protrusion-mediated Notch signaling.

**Figure 2.**
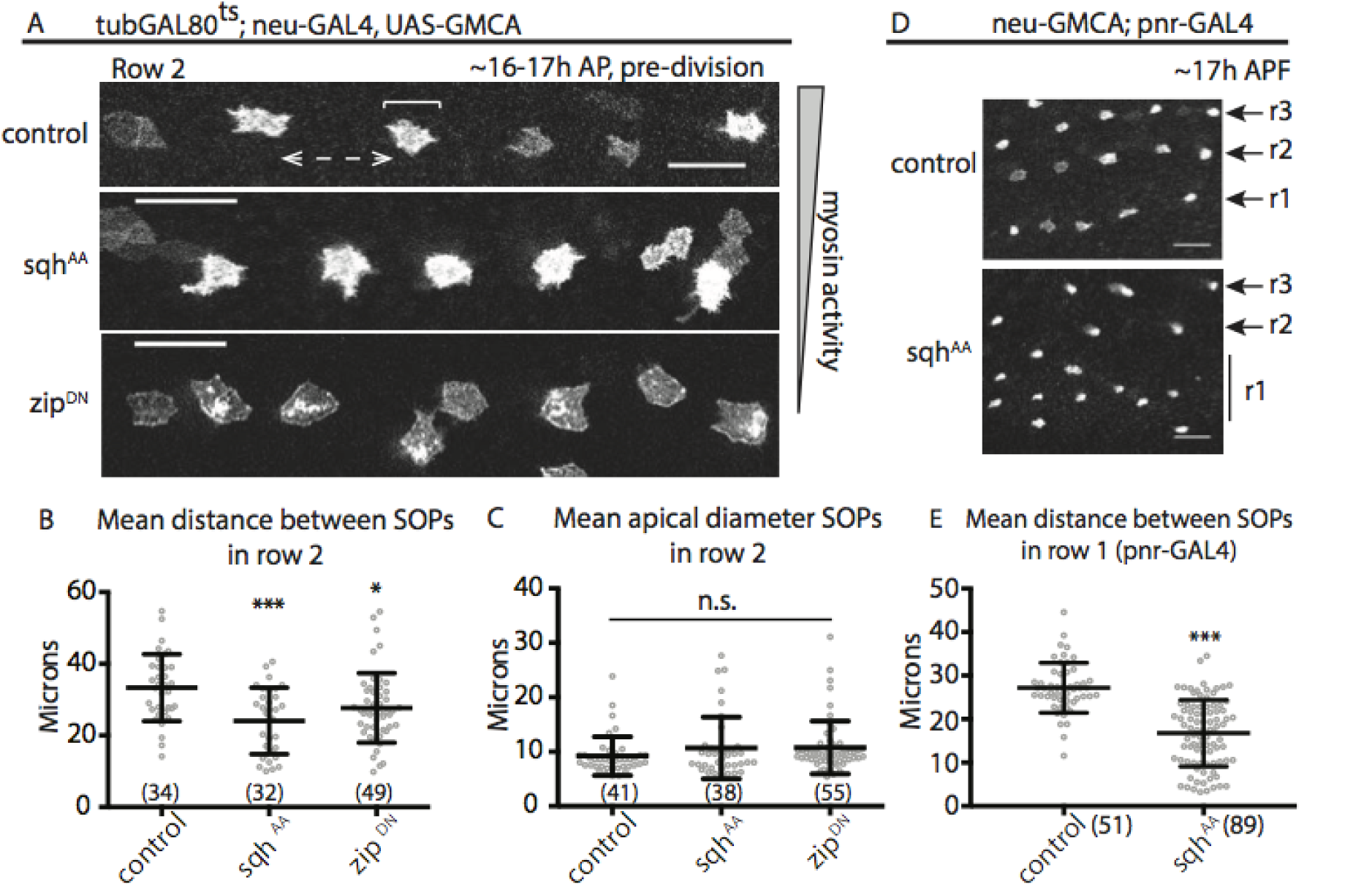
Changes in Myosin II activity disrupt bristle patterning. Expression of phospho-insensitive MRLC (sqh^AA^) or dominant negative zipper (zip^DN^) constructs in (A) SOP cells only disrupt the final SOP pattern in the notum. Row 2 is shown. (B) Mean **±** SD distance between SOPs in row 2 for the indicated genotypes (dashed, double-headed arrow in control (A)). (C) Mean **±** SD apical diameter for SOPs in row 2 for the indicated genotypes (see bracket in control (A)). (D) Expression of sqh^AA^ in all notum epithelial cells also disrupts the final SOP pattern in the notum. Rows are indicated by R, to the right of each panel. (E) Mean **±** SD distance between SOPs in row 1 for the genotypes in (D). Scale bars = 25*μ*m in all panels. ***, p ≤ 0.001; *, p ≤ 0.05; ns = not significant by one-way ANOVA with multiple comparisons. (n) = number of spaces (B) or cell diameters (C) measured. N ≥ 4 nota analyzed each genotype.

### The role of actomyosin contractility in the Notch signaling pathway

These data lead us to ask how actomyosin contractility contributes to the activity of the Notch signaling pathway. First, it is possible that non-muscle Myosin II activity is required for the Notch receptor and/or Delta ligand to be presented on the surface of signal sending and receiving cells (Koster et al., 2016). However, when we visualized Notch receptor and Delta ligand in SOP cells expressing the non-phosphorylatable Myosin II construct, we did not observe changes in the localization of Notch (apical or basal puncta) or Delta (cytoplasmic or basal puncta) (Figure S2). Notch and Delta were observed along protrusions regardless of Myosin II status (n >6 nota each genotype; Figure 3A-C). We conclude that lowering Myosin II activity does not affect the localization of Notch receptor or Delta ligand.

**Figure 3.**
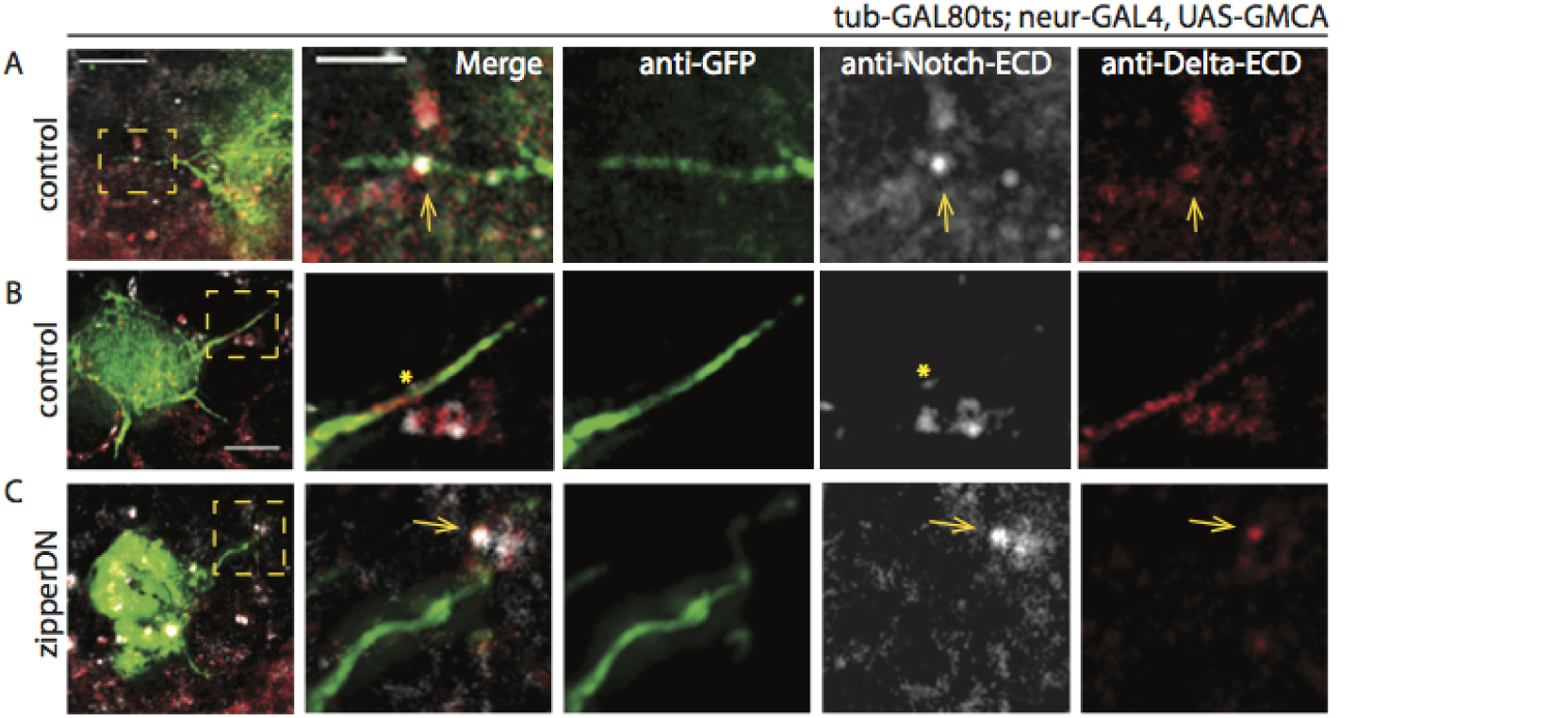
Localization of Notch and Delta. Fixed nota of the indicated GAL80, GAL4 genotypes crossed to (A-B) UAS-LifeActRuby or (C) UAS-zipperDN and co-immunostained for GFP (reporting filamentous Actin), Notch extracellular domain, and Delta extracellular domain. Scale bar, 5 *µ*m and 2 *µ*m. See also Figure S2.

Next, we asked whether Myosin II activity has an impact on basal protrusion morphology or dynamics. This is important, since previous genetic and mathematical modeling data show that decreased length or dynamics (i.e., retraction, extension) of basal protrusions can lead to defects in long-range Notch signaling, and therefore decreased SOP cell spacing (Cohen et al., 2010; Hadjivasiliou et al., 2016). We used time-lapse confocal microscopy to track the dynamics of basal protrusions in live SOP cells (Movie S1) marked using cell-type specific expression of the filamentous actin reporter GMCA, the GFP-tagged actin binding domain of Moesin (Kiehart et al., 2000)(Figure 4). Basal protrusions exhibited cycles of extension and retraction with a period of 10 minutes, with an average maximum length of 10 *µ* m, consistent with previously published data (Cohen et al., 2010). Protrusions changed shape prior to and during retraction (Movie S1), through a process reminiscent of helical buckling observed in mechanically-active filopodia in cell culture (Leijnse et al., 2015); this was observed in both control and sqh^AA^ expressing tissues (Movie S2). Moreover, expression of the squash^AA^ construct did not have a marked impact on protrusion morphology or dynamics compared to controls (Figure 4A-C). By contrast, the strong loss of Myosin II activity induced via expression of the dominant negative construct leads to aberrant protrusion morphologies (e.g., bulbous tips, fanning) and shorter protrusions (Figure 4D-D’), similar to the effect observed following Myosin inhibition in cell culture (Elliott et al., 2015). We conclude that while decreased Myosin II activity can lead to protrusion defects, these defects are unlikely to be sufficient to explain the reduction in protrusion-mediated long-range Notch signaling that we observed in animals expressing the squash^AA^ construct.

**Figure 4.**
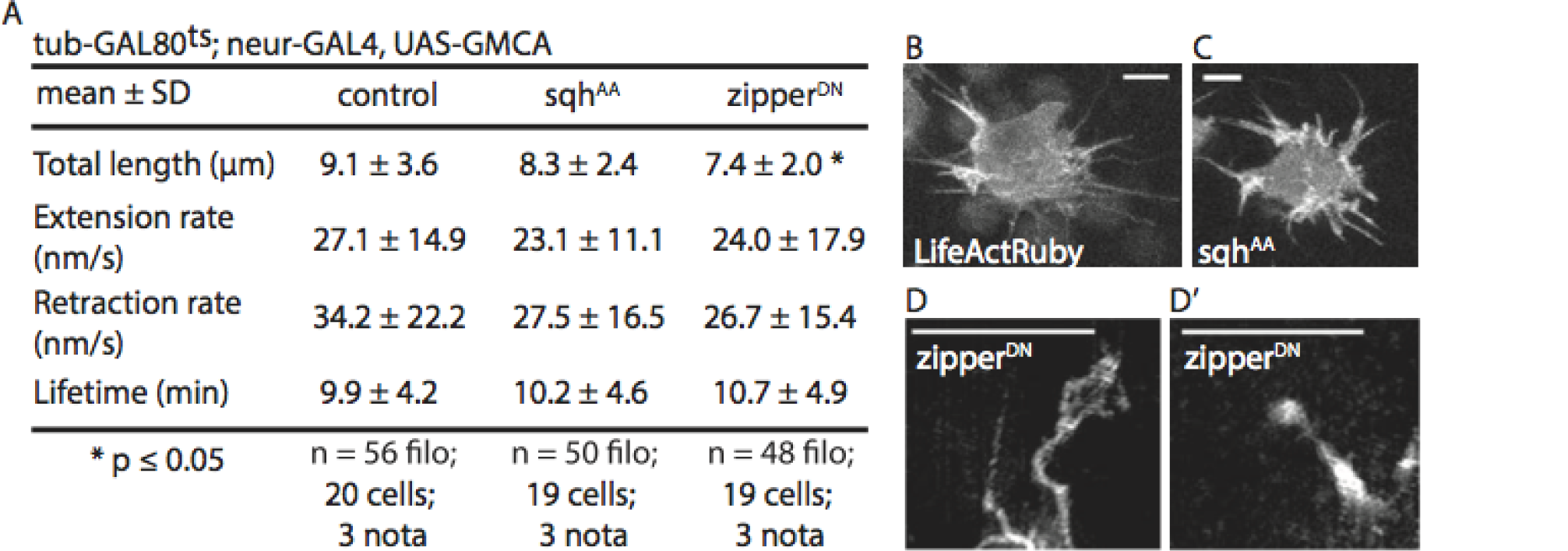
Effect of decreased NMM2 activity on protrusion dynamics and morphology. (A) For the listed genotypes, we measured maximum protrusion length (from cell body to tip), rate of extension (time to maximum length), rate of retraction (time to disappearance into cell body), and protrusion lifetime. (B-D’) Live imaging of SOP cells in nota of the indicated genotype show that overexpressing sqhAA does not appear to affect basal protrusions morphology, but overexpression of zipperDN does lead to abnormal protrusion shapes. Scale bars, 5*µ*m. *, p ≤ 0.05, by unpaired t-test. See also Supplemental Movies 1 and 2.

### Actomyosin contractility and endocytosis have overlapping roles towards Notch signaling

How might the actomyosin cortex influence Notch signaling? Endocytosis is an actin-dependent process that involves precise deformation of the cell cortex, whose stiffness is regulated by actomyosin contractility (Boulant et al., 2011; Liu et al., 2010). Moreover, the forces generated during ligand internalization have been proposed to drive the mechanical activation of Notch receptor (Gordon et al., 2015; Langridge and Struhl, 2017; Meloty-Kapella et al., 2012). This made it important to test whether the impact of actomyosin contractility on signaling and on lateral-inhibition patterning in vivo functions through its effects on endocytosis. We therefore used dsRNA to silence known regulators of Delta endocytosis and Myosin II activity to dissect their relationship in long-range Notch signaling (Figure 5A-B; S3). Decreased Delta endocytosis in SOP cells, induced by RNAi-mediated silencing of the epsin Liquid Facets (lqf) (Wang and Struhl, 2004), led to a failure to signal at adjacent contacts, and therefore clusters of SOP cells (10.7 ± 3.8 SOP pairs per LqfRNAi nota vs 4.5 ± 1.8 SOP pairs per control nota, N ≥ 3 nota each genotype; Figure S3B), as expected for a positive regulator of Notch signaling (Schweisguth, 2004). Next, to determine the relative contributions of actomyosin contractility and endocytosis to signaling, we combined this perturbation with Myosin RNAi. While the animals expressing dsRNAs that target each system alone had weak bristle patterning defects in this genetic background (Figure 5A), when we co-expressed zipper- and lqf-RNAi in SOP cells (using tubulin-GAL80^ts^ to temporally control RNAi expression so that the animals remained viable), we observed severe patterning defects. These appeared to be a combination of the two phenotypes observed in the single mutants, i.e., there was an increase in the variability of bristle spacing together with an enhanced number of GFP-positive cell clusters (Figure 5A-B). These data support the idea that Delta ligand endocytosis and Myosin II activity act in distinct ways to impact Notch signaling in vivo via lateral and protrusion-mediated signaling, respectively.

**Figure 5.**
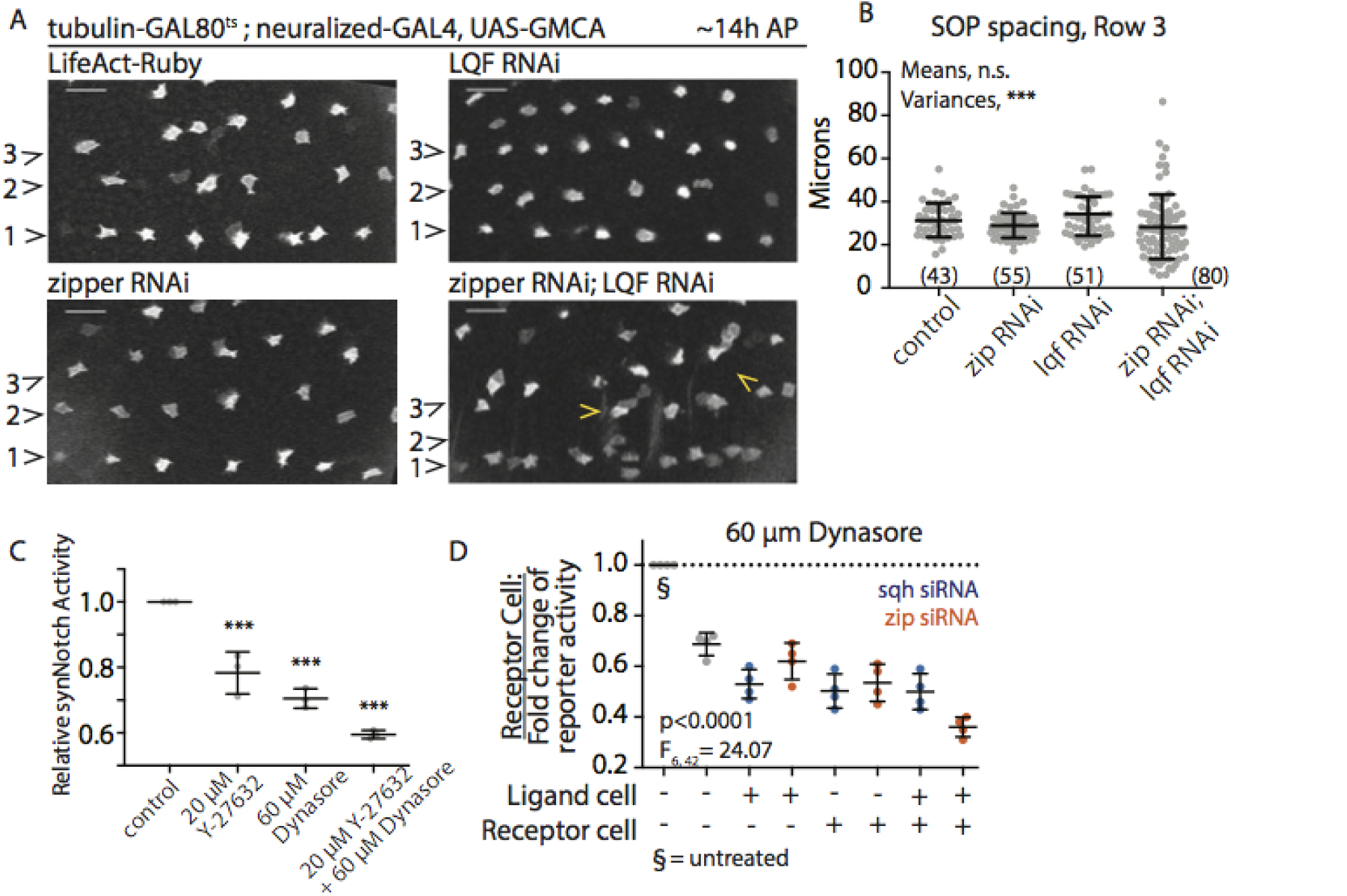
Overlapping roles of endocytosis and Myosin II activity for Notch activation. (A) SOP cell patterns for pupae of the indicated genotypes. Bristle rows 1–3 shown (indicated to the left of each panel). Scale bars, 25*μ*m. (B) Mean ± SD distance between SOP cells along bristle row 3 at 14 h AP for genotypes in (A); difference not significant for all comparisons. (n) = pairs measured, N ≥ 3 pupae per genotype. n.s. by one-way ANOVA. ***, p<0.0001 by Brown-Forsythe test, F3, 225 = 9.71. (C) synNotch activation in S2R+ heterogeneous populations treated with y-27632 alone, dynasore alone, or y-27632 and dynasore together. Mean ± SEM shown, ***, p≤ 0.001 by ANOVA with multiple comparisons. (D) Relative levels of synNotch activation in response to siRNA against zipper (orange) or squash (blue) in the presence of 60 *μ*M dynasore. + indicates transfection with siRNA, – indicates transfection with control siRNA targeting the white gene. Mean ± SEM for 4 experimental repeats shown. P-values determined by two-way ANOVA. Interaction term is not significant (F6,42 = 1.62). See also Figure S3.

This result was supported by experiments performed in cell culture using the synthetic Notch system, where we perturbed both Myosin II activity and endocytosis via dsRNA and pharmacological inhibition (Figure 5C-D). As previously published, Dynasore, an inhibitor of dynamin-mediated endocytosis (Macia et al., 2006), decreases Notch activation in this system (Gordon et al., 2015). Strikingly, when Dynasore was used combination with Y-27632, we observed a further reduction in Notch response (Figure 5C). Moreover, additive effects were observed in experiments in which we used dsRNAs to target either zipper or squash expression in signal sending or receiving cells in the presence or absence of Dynasore (Figure 5D). Again, these data point to Myosin II activity and endocytosis playing parallel roles in Notch signaling.

### Interacting protrusions are under mechanical tension

Since Myosin II appears to act in concert with endocytosis in Notch signaling, it could function to alter lateral inhibition through the generation of mechanical forces between basal, actin-based protrusions. In fact, in many cell types, Myosin II has been shown to drive both the flow of material from the tip to the base of filopodia towards the cell body (Lehmann et al., 2005) and filopodial retraction (Bornschlogl et al., 2013; Chan and Odde, 2008; Kress et al., 2007; Leijnse et al., 2015; Sayyad et al., 2015), while also generating tension in the cortical F-actin network that underlies the plasma membrane (Elliott et al., 2015; Fischer et al., 2009). Because of this, we considered the possibility that protrusions that make adhesive contacts with one another could exert forces on one another as they move, which could contribute to the activation of Notch receptor.

To test whether or not this is the case, we sought to establish whether Myosin II is present in the protrusions and then to determine if it acts locally to alter the mechanics of protrusions and the protrusion network as a whole. When we imaged Myosin II in the basal plane of the tissue, we observed phosphorylated, active, endogenous Myosin II localized in clumps at the base of protrusions in SOP cells (Figure 6A-A”). In addition, Venus-labelled Myosin II heavy chain (Lowe et al., 2014) was visible along the length of basal protrusions, where it co-localized with filamentous actin (Figure 6B-B”). These data are consistent with filopodial pulling and with models in which the contractile actomyosin meshwork at the base of cellular protrusions contributes to retrograde flow, amplifying actin treadmilling within the protrusion (Craig et al., 2012; Medeiros et al., 2006).

**Figure 6.**
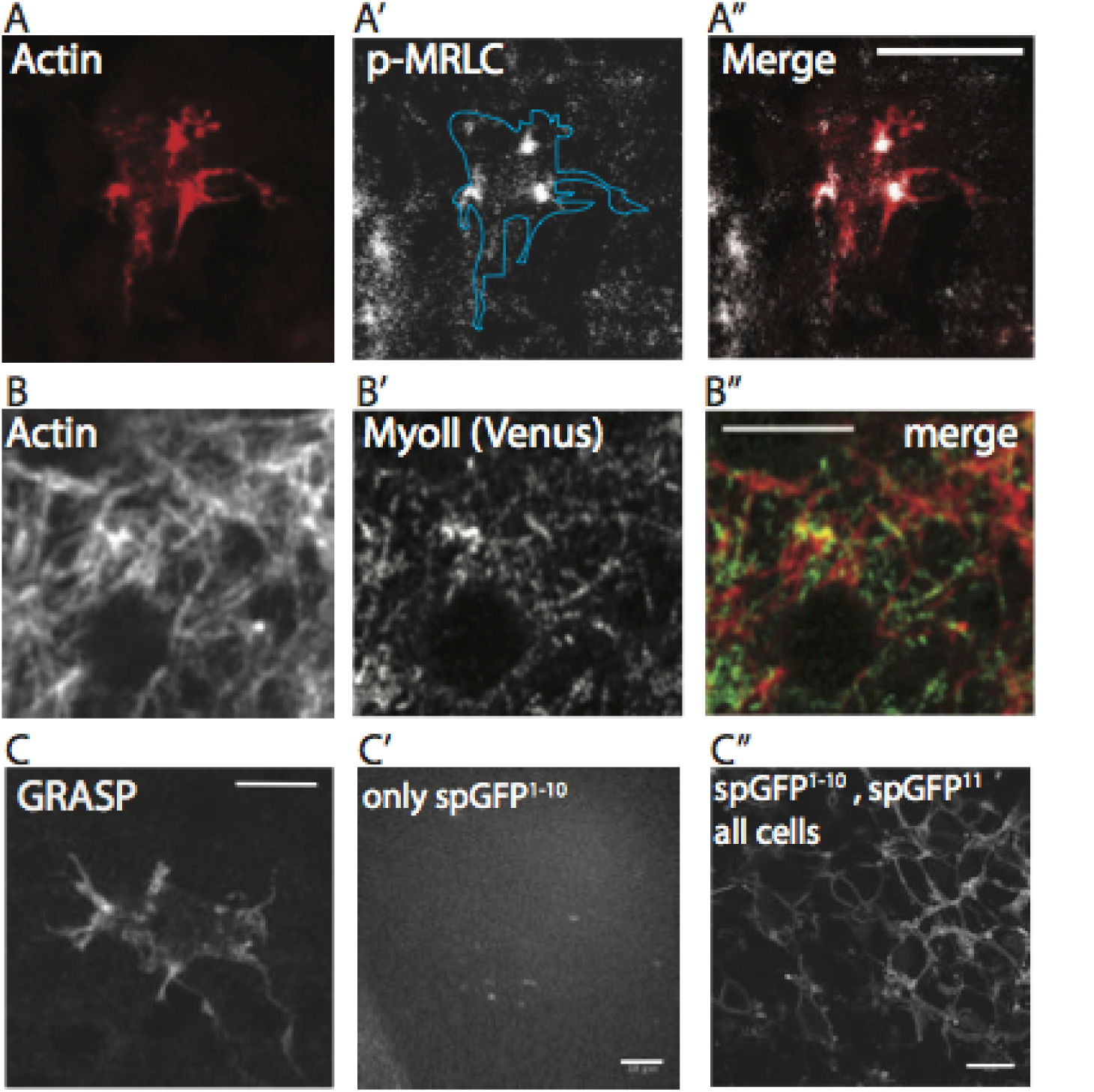
Protrusion contacts and Myosin II localization. (A-A”) Phosphorylated MRLC (squash) localizes to the cell-proximal base of basal protrusions in fixed nota. (B-B”) Filamentous actin in the basal protrusions co-colocalizes with Myosin II heavy chain (visualized with Venus) in live pupae. (C-C”) Image from a fixed nota of the indicated genotype, GRASP (reconstituted membrane bound GFP) indicates the extent of cell-cell contacts between signal sending (pictured) and receiving cells. Scale bars, 5 *µ*m, except in C’-C”, 10 *µ*m.

Having established the presence of Myosin II in protrusions, we wanted to assess the extent of contacts between the basal domain of cells in the notum. These can be visualized using the GRASP system (GFP reconstitution across synaptic partners)(Feinberg et al., 2008). Membrane-bound GFP^1-10^ was expressed in signal receiving cells and GFP^11^ in signal sending cells, so that the reconstituted GFP fluorescence would be visible when the two cell types came into contact with each other (Figure 6C). In tissues expressing only GFP^1-10^, we observe only background levels of fluorescence; in tissues where all cells express both GFP^1-10^ and GFP^11^ we observe all cell membranes fluoresce, including the nuclear envelope (Figure 6C’-C”). This revealed that basal cell-cell contacts are not restricted to protrusion tips, but are extensive and run along the length of the protrusions. As a result, protrusions form a network of basal cell-cell contacts that effectively extend the range of physical contact between cells in the notum to 2–3 cell diameters, as suggested previously (Cohen et al., 2010).

In line with this, when we imaged the full set of protrusions in vivo using pannier-GAL4 to express UAS-LifeActRuby in both signal sending and receiving cells in the notum, we observed a complex network of protrusions that crossed the basal surface of the epithelium (Figure 7A). These networks of fluctuate in density over time (Movie S3). During the window of observation, Particle Image Velocimetry (PIV)(Tseng et al., 2012) revealed that the basal domain is in constant flux (Figure 7A-B), without exhibiting any tissue-wide polarity. Importantly, this movement was dependent on actomyosin contractility, since it was markedly reduced by treatment with Y-27632 (Amano et al., 1996). Acute treatment with Y-27632 led to a steady decrease in the pulsatile movement of basal cell-cell contacts over a period of a few minutes (Figure 7C-D). These data suggest that basal contacts transmit forces between cells, which depend on Myosin II.

**Figure 7.**
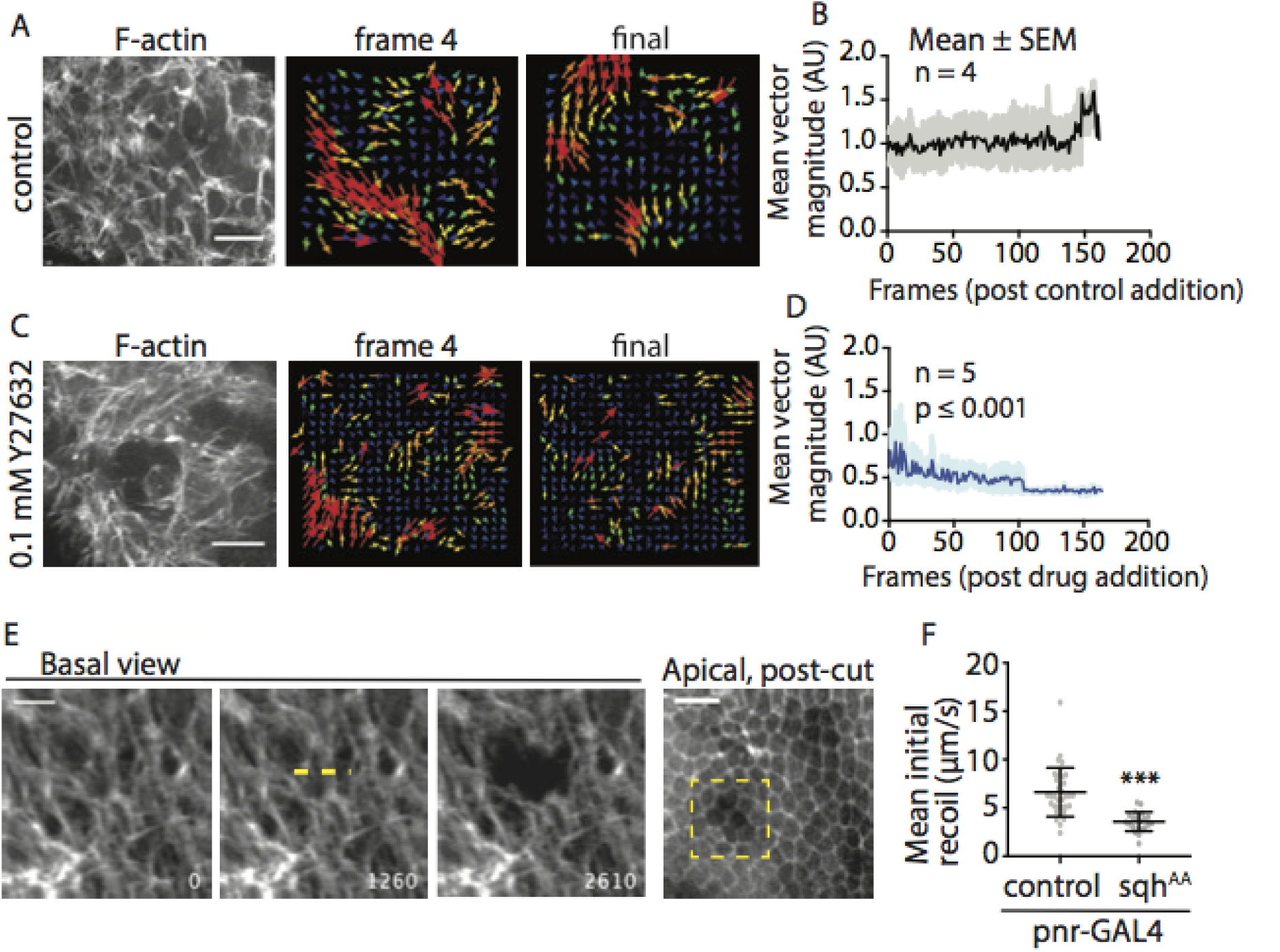
Basal protrusions and mechanical force. PIV analysis was performed after addition of either (A-B) control (dH2O) or (C-D) 0.1mM Y27632. (B, D) Quantification of mean vector magnitude for each frame pair after control or drug addition. Comparison of best-fit slope relative to control, *** p<0.0001. (E) Basal view of live nota expressing UAS-LifeActRuby under pnr-GAL4 in all signaling cells during a laser ablation protocol. Dashed yellow line indicates location and length of cut target. Time, in ms, at lower right. Apical view post-cut indicates no apparent whole cell ablation in the area where the basal surface was targeted (yellow box). Scale bars: basal view, 10 *µ*m; apical view, 25 *µ*m. (F) Quantification of mean initial recoil post-laser ablation of the basal protrusion network for control (UAS-LifeActRuby) or sqhAA expressing nota. ***, p≤ 0.001 by unpaired t-test. See also Supplemental Movie 3.

Finally, to test whether these basal protrusion contacts are under tension, we performed laser ablation to disrupt their connections. Using a focused UV-laser, we made 10 *µ*m cuts on the basal surface of the notum, disrupting the protrusion networks without ablation of the rest of the cell body in more apical planes (Figure 7E). We observed that the cuts induce a rapid recoil, followed by a slower relaxation, as expected for tissues under tension. When we expressed the sqh^AA^ construct under the pnr-GAL4 driver to reduce Myosin II activity throughout the notum, we observed a two-fold decrease in the mean initial recoil velocity, from 6.6 ± 2.5 *µ*m/s (mean ± SD, n = 35 cuts, 10 pupae; Figure 7F) to 3.6 ± 1.0 *µ*m/s (n = 22 cuts, 8 pupae; p <0.0001, unpaired t-test). Our ablation data show that protrusion contact networks exert mechanical forces on each other in a Myosin II dependent manner. Altogether, these data support a model of protrusion connection and contractility that requires localized Myosin II activity for the robust activation of Notch response during pattern formation in the notum.

## Discussion

In Drosophila, the pattern of bristles on the dorsal thorax of the fly is set up by Notch-Delta signaling through a process of lateral inhibition (Bray, 2016). As shown in our study, this involves two distinct processes. First, there is lateral inhibition between adjacent cells, which is enhanced by both actomyosin contractility and endocytosis (Musse et al., 2012). While a reduction in Myosin II activity in signal sending cells is not sufficient to induce adjacent cells to adopt the SOP cell fate, in combination with decreased ligand endocytosis, we observe frequent SOP cell clusters. This suggests that Notch signaling in adjacent neighbors depends on both types of force-dependent signaling. This makes sense, since cortical actomyosin is required for endocytosis to pull against when generating forces. Second, over longer distances, where lateral inhibition signaling relies on protrusions, reductions in Myosin II activity are sufficient to lead to aberrant cell fate decision making – leading to decreased spacing between SOP cells. This suggests a more important role for Myosin II during these types of signaling events.

How might Notch receptor be activated on protrusions? Our data suggests that one way this could occur is through the contact, engagement, and retraction of basal protrusions in Notch and Delta expressing cells. Importantly, since the contact area between protrusions can be extensive, Notch/Delta signaling molecules localized along the protrusions could potentially contribute to signal activation. In addition, once contact has been made, the ligand and receptor could diffuse along the length of the protrusion and become trapped at the site of contact – as previously suggested (Khait et al., 2016). Once Notch and Delta have become engaged, actomyosin-dependent extension/retraction cycles could provide shear forces parallel to the plasma membrane that contribute to the activation of Notch. Alternatively, Notch could be activated by the simple pulling of protrusions on one another. Development of highly sensitive Notch activity reporters, that allow the rapid visualization of local receptor activation in vivo will be necessary to reveal the mechanical details of Notch signaling in cellular protrusions.

Our data do not rule out the possibility that endocytosis contributes directly to protrusion-mediated Notch activation. As in the case of signaling between adjacent cells, actomyosin contractility may augment the physical effects of endocytosis by providing a rigid cell cortex against which pulling forces generated during the internalization of a clathrin-coated pit can act (Xie et al., 2012). However, the structure of filopodial-like protrusions, with bundled actin closely associated with the plasma membrane, may be refractory to endocytosis. Indeed there is evidence for the existence of endosomal ‘hot spots’ at the base of cellular protrusions rather than along them (Heusermann et al., 2016). A closer analysis of the ultrastructure of basal protrusions could help to reveal the impact of local structure on the numbers of endocytic pits formed.

How much does the protrusion-mediated Notch-Delta signaling contribute to the cell fate decision making process of epithelial cells? Previously, it was shown that stochastic noise can be a feature of signaling mechanisms and contribute to the plasticity of bristle patterning (Cohen et al., 2011). More generally, stochastic amplification of noisy signals, especially as part of signaling feedback loops, can drive bistable systems in both models and biological systems (Eldar and Elowitz, 2010; Hsu et al., 2016; Losick and Desplan, 2008; Palani and Sarkar, 2012). From our results, it is clear that the amount of signaling protein that is present on basal protrusions is sufficient to induce changes in cell fate: when two cells beginning to downregulate Notch response and upregulate pro-neural genes are “too close” i.e., within range of each other’s basal protrusions, we commonly observe switching away from the SOP fate in at least one of them. Thus, the long-range Delta signal, perhaps amplified through the feedback mechanisms which exist to reinforce Notch-mediated cell fate decisions (Heitzler et al., 1996; Parks et al., 2008), is sufficient to impose the epithelial fate, even if these proteins are found concentrated at lateral cell-cell contacts (Cohen et al., 2010; Trylinski et al., 2017)

Recent evidence shows that the patterning of bristle rows begins developing in the notum far earlier than 12 h APF, where we begin our analysis (Corson et al., 2017). Although we have observed basal protrusions in fixed nota as early as 9hr APF (unpublished data), technical challenges prevent us from being able to perform our live analysis of Notch dynamics prior to 12 h APF. However, our data clearly show that the SOP-epithelial fate decision is a temporally extended process; these fate choices remain plastic as late as 12 hours APF in wildtype animals (Doe and Goodman, 1985). However, given that Myosin II is able to alter Notch-Delta signaling in cell culture and in vivo, and given the prevalence of protrusions in biological systems (Kornberg, 2017), it will be important in the future to determine whether actomyosin-dependent protrusion-mediated Notch signaling plays much more general roles in tissue development and homeostasis than currently appreciated.

In summary, these data provide the first evidence that Myosin II helps generate the force necessary to activate Notch following ligand binding, and reveal a specific function for Myosin II in this type of signaling during protrusion-mediated lateral inhibition in the developing fly.

## Methods

### Fly strains

GAL4 Drivers: tubulin-GAL80^ts^; neuralized-GAL4, UAS-GMCA. Neur-GMCA; pnr-GAL4. NsfGFP; neur-GAL4. NsfGFP; pnr-GAL4. shotgun^GFP^; pnr-GAL4. shotgun^GFP^, neur-GMCA; neur-GAL4. UAS-lifeActRuby; pnr-GAL4. UAS Responders: UAS-zipper^dn, gfp^. UAS-squash^AA^. UAS-LifeActRuby. UAS-liquid facets RNAi. UAS-ROK RNAi. UAS-white RNAi UAS-zipper RNAi. Other: zipper^Venus, Flag^.

### Microscopy

White pre-pupae were picked and aged to 12-24 h AP at 18°C or 30°C (for GAL80^ts^ lines). Live pupae were removed from pupal case and mounted on a slide as previously described (Loubery and Gonzalez-Gaitan, 2014). Final patterns, live imaging of NsfGFP, ex vivo experiments, and filopodia imaging was performed on either a Leica SPE confocal, 40x oil immersion objective (1.15 NA) at room temperature or a Nikon EclipseTi, 20x (0.75 NA) or 60x (1.4 NA). Localization of myosin in live pupae was performed on a Zeiss LSM880 with AiryScan, 63x (1.4 NA) oil immersion objective. Fixed nota were imaged on a Leica SPE confocal, 63x (1.3 NA) oil immersion objective. Fixed SIM images were obtained on a Zeiss Elyra PS.1, 63x (1.4NA) oil immersion. Fixed Delta uptake images acquired on DeltaVision confocal, 100x (1.4 NA) oil immersion objective. Laser ablation: Live images were acquired on a Zeiss Axio Imager.M2m, 40x (1.2NA) water objective, using Micromanager (Vale Lab) acquisition software. An Nd:YAG UV laser (Continuum) was interfaced with the confocal microscope to allow steered laser incisions (Hutson et al., 2003). We made 10*µ*m incisions orthogonal to the anterior-posterior axis at a laser power 3.0*µ*J.

### Immunofluorescence

White pre-pupae were picked and aged to 12-24 h AP at 18°C or 30°C (for GAL80^ts^ lines). At specified age, the nota were dissected and fixed in 4% paraformaldehyde in 1X PBS (0.05% Tween). Tissues were blocked in 1X PBS (0.05% Tween) with 5% bovine serum albumin (Sigma) and 3% fetal bovine serum (Sigma). Primaries: chicken anti-GFP (1:1000, Abcam); rabbit anti-pS19-MRLC (1:50, CST); mouse anti-Delta extracellular domain (1:100, DSHB); mouse anti-Notch extracellular domain (1:100, DSHB); Guinea pig anti-Delta ECD (1:2000, M. Muskavitch); mouse anti-Notch ECD (C458.2H, 1:200, DSHB). Secondaries: AlexaFluor 488 anti-chicken and AlexaFluor 568 anti-rabbit (ThermoFisher Scientific, 1:1000); Rhodamine Red-X anti-guinea pig and Cy5 anti-mouse (Jackson ImmunoResearch Laboratories, both 1:2000). Texas Red Phalloidin (AlexaFluor, 1:500) was used to visualize F-actin. Nota mounted in 50% glycerol or Vectashield with DAPI (Vector Laboratories).

### Ex vivo

Live nota were dissected and attached to 35mm^2^ glass bottom dishes (Matek) using a thrombin/fibrinogen (Sigma) clot, then cultured in 250*µ*L modified Clone8 medium (Schneider’s insect medium (Sigma), 2.5% fly extract, 2% fetal bovine serum (ThermoFisher Scientific)) as previously described (Loubery and Gonzalez-Gaitan, 2014). After initiation of imaging, either control (250*µ*L Clone8 + 0.5*µ*L dH2O) or 0.1mM Y27632 (250*µ*L Clone8 + 0.5*µ*L 100mM Y27632; Sigma) was added to the dish.

### Molecular Cloning

Long hair-pin dsRNA was designed using SnapDragon (DRSC/TRiP Functional Genomics Resources, http://www.flyrnai.org/snapdragon). Gene-specific amplicons of sqh, zipper, and white genes were amplified from fly genome by PCR, inserted into pDONR221 vector, and subcloned into pWALIUM10 RNAi vector. All constructs verified by sequencing. The primers used for the synthesis of dsRNA were as follows (F, forward; R, reverse), sequence 5’-3’:

sqh F: GGGGACAAGTTTGTACAAAAAAGCAGGCTTCCGGCTCCATTTAGCTCCATTA;
sqh R: GGGGACCACTTTGTACAAGAAAGCTGGGTGGCAGGACGCCCATATTCTC;
zipper F: GGGGACAAGTTTGTACAAAAAAGCAGGCTTCCACAGGGTACAGCCGATAAA;
zipper R: GGGGACCACTTTGTACAAGAAAGCTGGGTGCTTGTGCTTGCAGCTTCTTC;
wF: GGGGACAAGTTTGTACAAAAAAGCAGGCTTCTGCCCAGTGTCCTACCA;
wR: GGGGACCACTTTGTACAAGAAAGCTGGGTGTACGAGGAGTGGTTCCTTGA.

Primers used for qPCR:

GAPDH F: CCAATGTCTCCGTTGTGGA
GAPDH R: TCGGTGTAGCCCAGGATT
sqh F1: CGAGGAGAATATGGGCGTCC
sqh R1: CCTCCCGATACATCTCGTCCA
zipper F: CCAAGACGGTCAAAAACGAT
zipper R: GATGTTGGCTCCCGAGATAA

qPCR: S2R+ cells plated in 6-well plate were transfected with indicated dsRNAi plasmid (0.6 *µ*g) together with pAct-GAl4 (0.2 *µ*g) using Effectene. dsRNA against white used as control. The transfection procedure was repeated two more times every 4 days to maximize transfection efficiency. Total RNA was extracted from S2R+ cells after the 3rd transfection using TRIzol Reagent (Thermo Fisher Scientific). Raw RNA was treated with DNase I, purified by QIAGEN RNeasy kit, and converted to cDNA template using iScript cDNA Synthesis Kit (Bio-Rad). Y27632 treatment: Cells expressing either synNotch or GFP-ligand were treated with 20*µ*M Y-27632 for 1 hour and then mixed and cultured in the presence of 20*µ*M Y-27632 for 1 day before analysis.

### Cell Culture

Drosophila S2R+ cells were grown in Schneider’s Drosophila media (Gibco) supplemented with 10% fetal bovine serum and 0.5% Pen/Strep (Gibco). Cells were transfected using Effectene transfection reagent (Qiagen) (Zhou et al., 2013), with DNA mixture containing: 0.1 *µ*g pAct-Gal4, 0.4 *µ*g pUAST-dsRNA, 0.01*µ*g pUbi::GBN-Notch-QF, 0.09 *µ*g pQUAST-luciferase (for signal receiving cells); 0.1*µ*g pAct-Gal4, 0.4 *µ*g pUAST-dsRNA, 0.1 *µ*g pUbi::GFP-mcd8-Ser (for signal sending cells). Transfected cells were cultured for 10 days to allow efficient knock-down of target genes. Signal sending and receiving cells were washed twice with fresh culture medium, suspended, and mixed together in 1:1 ratio. Cells were cultured for an additional day before screening luciferase activity (Steady-Glo Luciferase Assay Kit, Promega) using a SpectraMax Paradigm Multi-Mode Microplate Reader. For Dynasore treatment, transfected cells were washed twice by culture medium, incubated with medium containing 60*µ*M Dynasore and/or 20*µ*M Y-27632 (final concentration) for 1 hr. Cells were resuspended, mixed in 1:1 ratio, and cultured for an additional day in the presence of Dynasore before luciferase assay. Knock-down efficiency was tested by qPCR. Real-time PCR was performed using iTaq SYBR Green Supermix (Bio-Rad) with GAPDH as a control.

### Quantitative analysis

Figure 1: NsfGFP signal was measured as previously described (Hunter et al., 2016).

Figure 2, 5: Mean distance and apical diameter between SOPs was measured in bristle row 2: an unprojected, z-slice image is rotated to align bristle row 2 with 0° axis, an ROI is drawn to encompass all SOPs in row 2, the ROI is collapsed along the y-axis to result in a 1D histogram along the bristle row (x-axis). Distance between SOPs is defined as the centroid to centroid distance between signal peaks along the histogram; apical diameter is defined as the edge-to-edge measurement of a signal peak (i.e., where signal reaches 0).

Figure 4: Live images were maximum projected. Total length = maximum length from cell-proximal base to tip; Extension rate = time for a protrusion to appear until it reaches maximum length; Retraction rate = time for a protrusion to disappear after reaching maximum length; Lifetime: total time during which a protrusion is visible.

Figure 7: Particle Image Velocimetry (PIV) was performed using the FIJI plugin described in (Tseng et al., 2012). Mean vector magnitude (second iteration) for the field of view for each timepoint pair was calculated. Mean initial recoil is measured by the average movement/time of ≥ 2 fiducials before and after laser ablation, and was performed using the manual spot tracker in Icy (de Chaumont et al., 2012).

Supplement: Filopodia dynamics were measured by manually tracing protrusions in z-projected time-lapse images. Myosin localization: basal images (up to 1*µ*m maximum projection). ROIs were drawn to include only basal filopodia areas.

## Acknowledgements

We thank Dan Kiehart (Duke University) for UAS-zipper^DN, GFP^ stocks and Marc Muskavitch for Delta antibody (guinea pig). Fly stocks obtained from the Bloomington Drosophila Stock Center (NIH P40OD018537) were used in this study. Laser ablation experiments were performed with the help of the Kiehart Lab (Duke University). Microscopy was performed with the help of Vincent Schram (NICHD) and Carolyn Smith (NINDS). We thank Dr. Stephane Romero for his critical reading and comments. We are thankful to members of the Baum, Charras, and Giniger Labs for their technical advice and manuscript edits. Project funding was acquired by G.C., E.G., and B.B. Overall direction of the project by G.H., E.G., G.C., and B.B. This work was supported by the National Institutes of Health (Z01 NS003013 to E.G. and R21DA039582 to N.P.); the Howard Hughes Medical Institute (N.P.); a Daymon Runyon Cancer Research Foundation (to L.H.) the Biotechnology and Biological Research Council (BB/J008532/1 to B.B. and G.C.); a Cancer Research UK fellowship (to B.B.) and University College London (to B.B.). Laser ablation microscopy in the Kiehart lab is supported by NIH GM033830 to DPK.

## Author contributions

G.L.H., L.H., E.G., and B.B. wrote the manuscript. G.L.H. and B.B. designed experiments. G.L.H. performed and analyzed fly experiments, L.H. performed and analyzed cell culture experiments.

## Supplemental Figures

Movie S1. Basal protrusion dynamics in a SOP cell expressing UAS-GMCA under the neur-GAL4 driver. Movie represents a maximum projection over 2.1 *µ*m. Time in seconds. Scale bar, 10 *µ*m.

Movie S2. Basal protrusion dynamics in a SOP cell expressing UAS-GMCA and UAS-sqhAA under the neur-GAL4 driver. Movie represents a maximum projection over 2.1 *µ*m. Time in seconds. Scale bar, 10 *µ*m.

Movie S3. Basal protrusion dynamics in cells expressing UAS-LifeAct-Ruby under the pnr-GAL4 driver to visualize filamentous actin in all epithelial cells. This movie is from a tissue explant cultured in Clone8 medium. Movie represents a single z-plane, time stamp: (minutes:seconds). Scale bar, 10 *µ*m.

**Figure S1.**
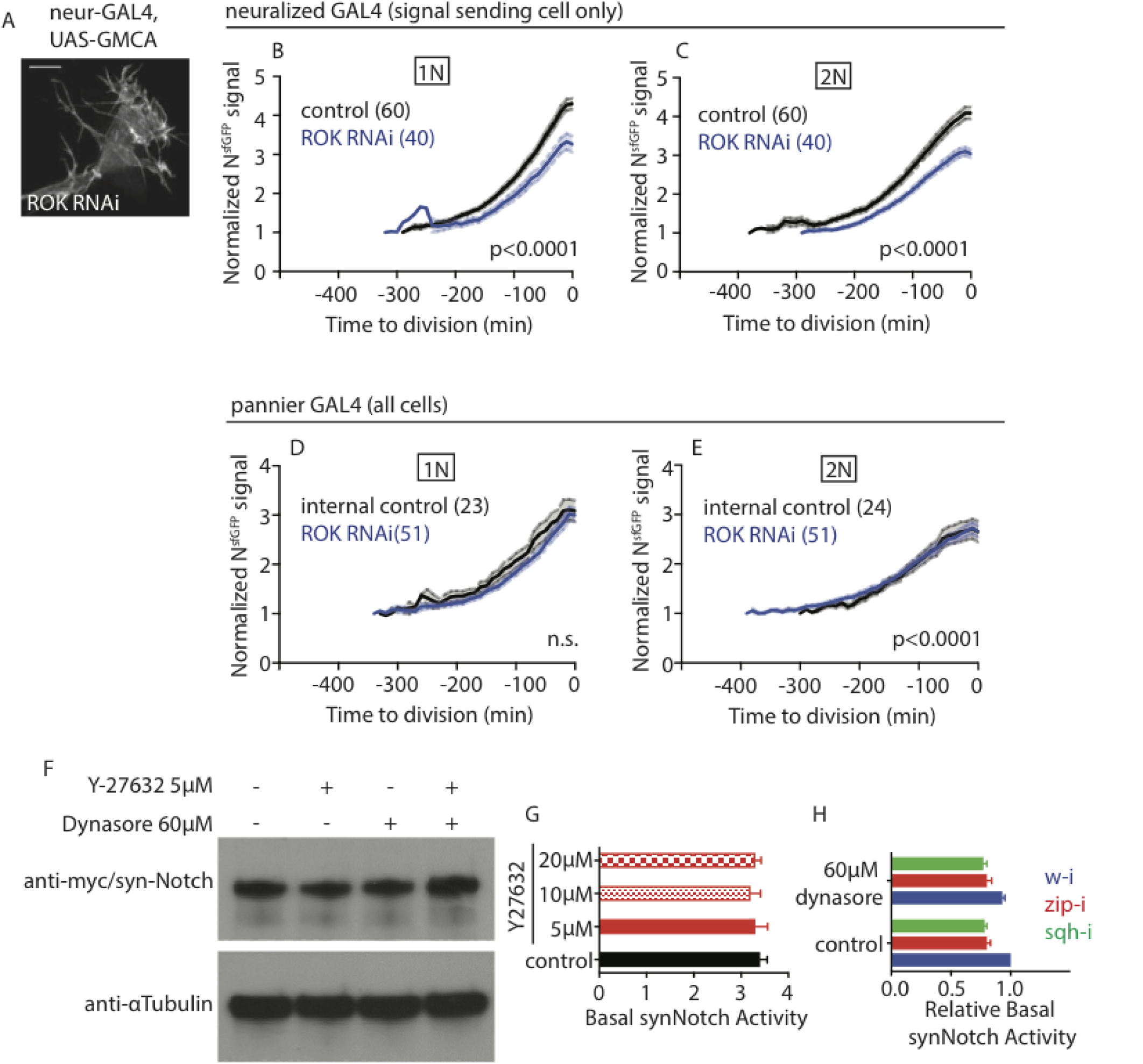
Contribution of Myosin II activity to Notch response. (A) RNAi against an activator of Myosin II activity, Rho kinase (ROK) in signal sending cells alone does not disrupt protrusion morphology. (B-C) Decreased ROK activity in signal sending cells alone (via neur-GAL4) leads to decreased Notch response in both (B) adjacent and (C) distant wildtype neighboring cells (non-linear regression, comparison of fit, Prism). (D) Decreased ROK activity in all cells (via pnr-GAL4) does not affect the rate of signaling between adjacent cells, but does decrease the total signal (1N control vs RNAi elevations, p¡0.001 by linear regression). (E) The rate of Notch response in distant neighbors is significantly affected by ROK RNAi expression (linear regression, Prism). (F) S2R+ cells expressing synNotch in the absence of ligand expressing cells and cultured in the presence of Y27632 and/or Dynasore do not exhibit changes in their expression of synNotch in response to drug treatment. (G) Acute inhibition of ROK does not significantly alter the basal synNotch activity measured in S2R+ cells expressing synNotch in the absence of ligand expressing cells. (H) Basal synNotch activity is affected by transfection of zip and sqh siRNA, but not by acute treatement with Dynasore. However, because fold changes of synNotch activity in the presence of GFP-ligand is calculated based on each treatment respectively, the relative fold changes should still primarily reflect the efficiency of synNotch cleavage under different conditions.

**Figure S2.**
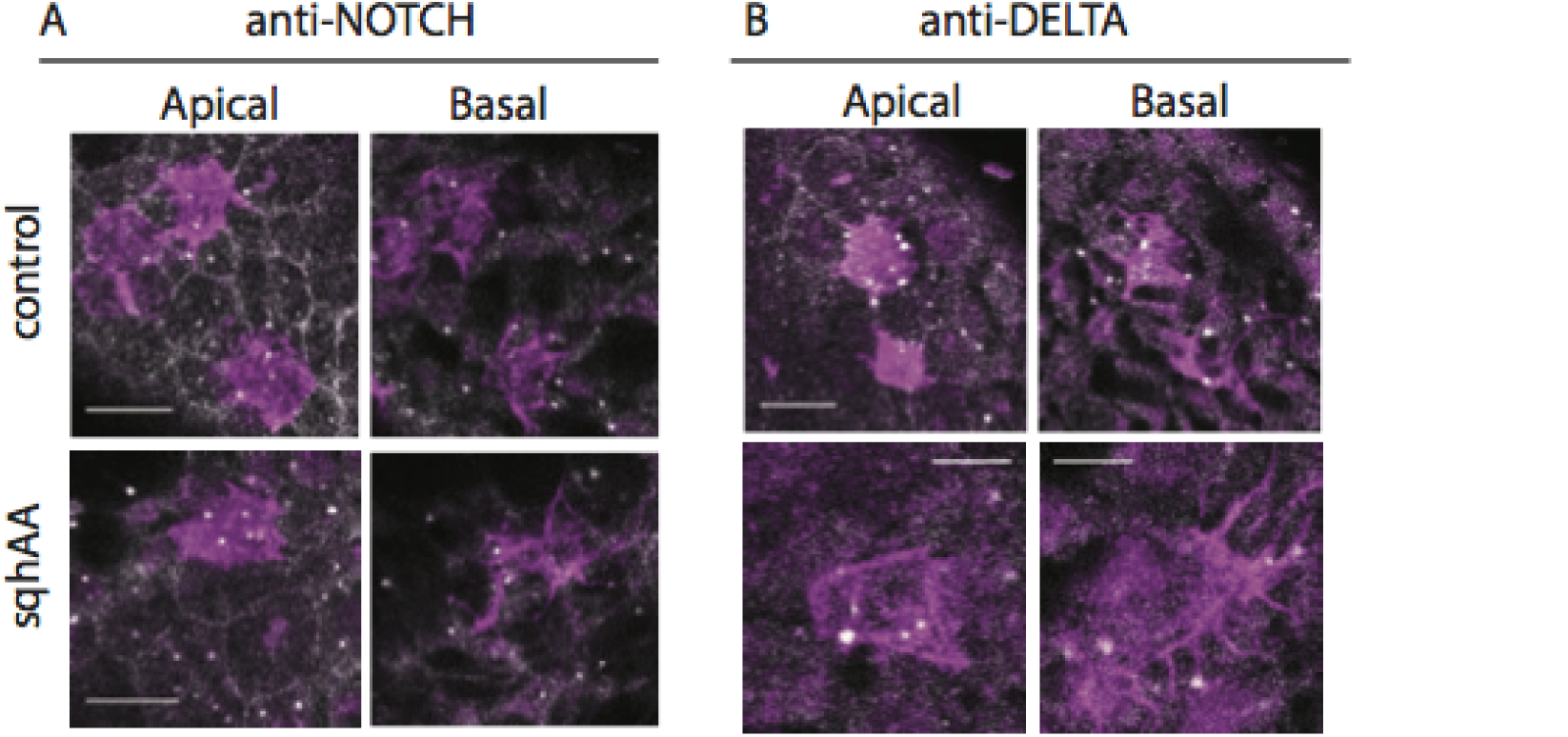
Localization of Notch and Delta with decreased Myosin II activity. In tissues expressing control (LifeActRuby) or sqh^AA^ constructs in SOP cells (tubGAL80^ts^; neur-GAL4, UAS-GMCA) we observe no differences in (A) Notch localization or (B) Delta localization at a single apical section and basal projection (over 2 microns). Scale bars, 10 *µ*m and 5 *µ*m (sqh^AA^ anti-Delta panels).

**Figure S3.**
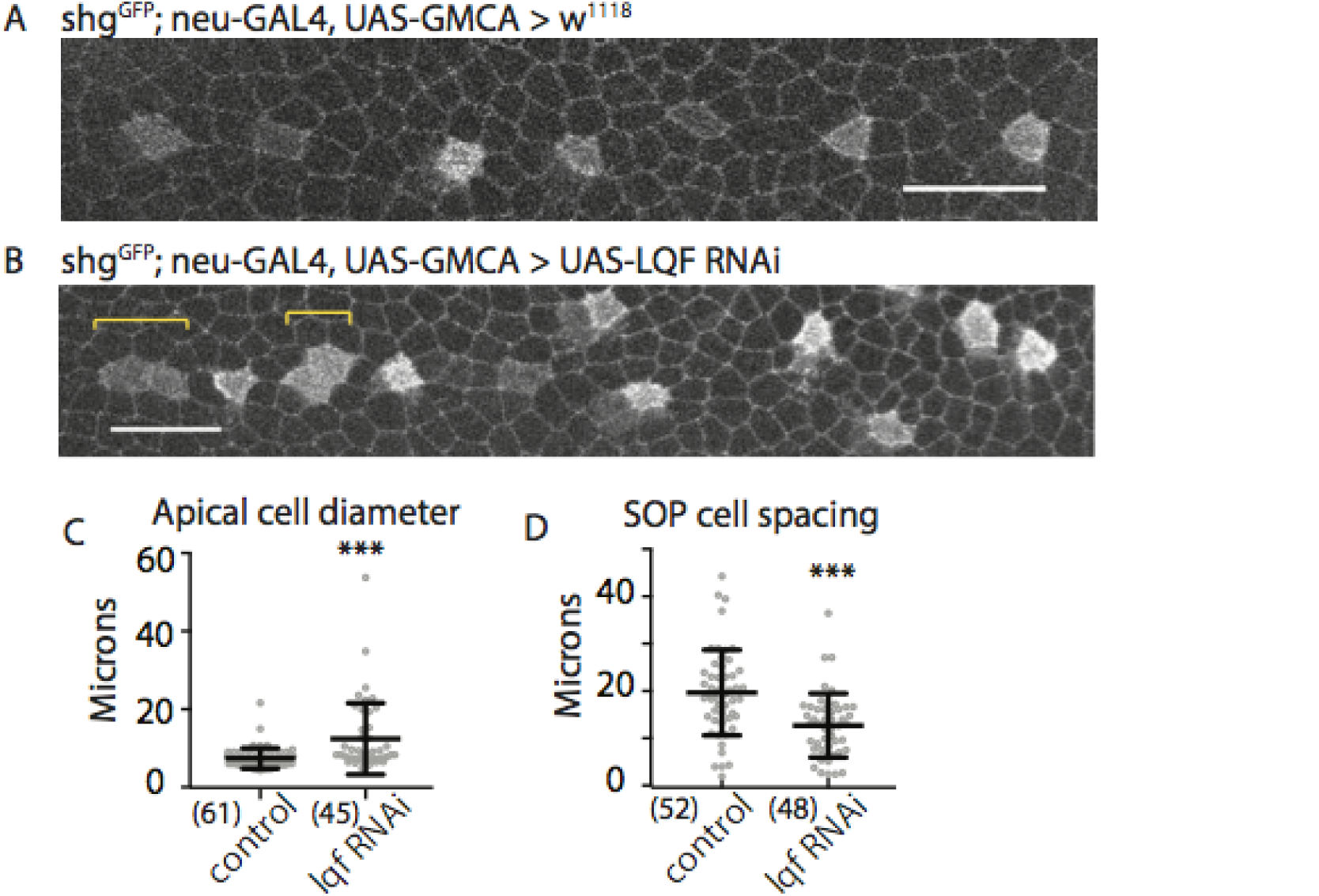
Role of ligand endocytosis in Notch signaling. (A-B) RNAi against key regulator of ligand endocytosis pathway (liquidfacets) lead to a pairing phenotype (indicated by yellow brackets). Pairs of neuralized expressing cells are seen with less frequency in control pupae. (A-B) Scale bars, 25 *µ*m. (C-D) Quantifications of patterns for genotypes in (A-B). (C) Apical ‘cell’ diameter, measuring grouping/pairing. ***, p <0.001 by unpaired t-test. (D) Measurements of the distance between neuralized expressing cells. ***, p <0.001 by unpaired t-test. (n), number of distances measured, N ≥ 3 nota measured for each genotype at 14 h AP.

## References

Amano, M., Ito, M., Kimura, K., Fukata, Y., Chihara, K., Nakano, T., Matsuura, Y., and Kaibuchi, K. (1996). Phosphorylation and activation of myosin by Rho-associated kinase (Rho-kinase). J Biol Chem 271, 20246–20249.

Artavanis-Tsakonas, S., and Simpson, P. (1991). Choosing a cell fate: a view from the Notch locus. Trends Genet 7, 403–408.

Basu, S., and Proweller, A. (2016). Autoregulatory Control of Smooth Muscle Myosin Light Chain Kinase Promoter by Notch Signaling. J Biol Chem 291, 2988–2999.

Bischoff, M., Gradilla, A.C., Seijo, I., Andres, G., Rodriguez-Navas, C., Gonzalez-Mendez, L., and Guerrero, I. (2013). Cytonemes are required for the establishment of a normal Hedgehog morphogen gradient in Drosophila epithelia. Nat Cell Biol 15, 1269–1281.

Bornschlogl, T. (2013). How filopodia pull: what we know about the mechanics and dynamics of filopodia. Cytoskeleton (Hoboken) 70, 590–603.

Bornschlogl, T., Romero, S., Vestergaard, C.L., Joanny, J.F., Van Nhieu, G.T., and Bassereau, P. (2013). Filopodial retraction force is generated by cortical actin dynamics and controlled by reversible tethering at the tip. Proc Natl Acad Sci U S A 110, 18928–18933.

Boulant, S., Kural, C., Zeeh, J.C., Ubelmann, F., and Kirchhausen, T. (2011). Actin dynamics counteract membrane tension during clathrin-mediated endocytosis. Nat Cell Biol 13, 1124–1131.

Bray, S.J. (2016). Notch signalling in context. Nat Rev Mol Cell Biol 17, 722–735.

Chan, C.E., and Odde, D.J. (2008). Traction dynamics of filopodia on compliant substrates. Science 322, 1687–1691.

Cohen, M., Baum, B., and Miodownik, M. (2011). The importance of structured noise in the generation of self-organizing tissue patterns through contact-mediated cell-cell signalling. J R Soc Interface 8, 787–798.

Cohen, M., Georgiou, M., Stevenson, N.L., Miodownik, M., and Baum, B. (2010). Dynamic filopodia transmit intermittent Delta-Notch signaling to drive pattern refinement during lateral inhibition. Dev Cell 19, 78–89.

Corson, F., Couturier, L., Rouault, H., Mazouni, K., and Schweisguth, F. (2017). Self-organized Notch dynamics generate stereotyped sensory organ patterns in Drosophila. Science 356.

Craig, E.M., Van Goor, D., Forscher, P., and Mogilner, A. (2012). Membrane tension, myosin force, and actin turnover maintain actin treadmill in the nerve growth cone. Biophys J 102, 1503–1513.

Curran, S., Strandkvist, C., Bathmann, J., de Gennes, M., Kabla, A., Salbreux, G., and Baum, B. (2017). Myosin II Controls Junction Fluctuations to Guide Epithelial Tissue Ordering. Dev Cell.

de Chaumont, F., Dallongeville, S., Chenouard, N., Herve, N., Pop, S., Provoost, T., Meas-Yedid, V., Pankajakshan, P., Lecomte, T., Le Montagner, Y., et al. (2012). Icy: an open bioimage informatics platform for extended reproducible research. Nat Methods 9, 690–696.

De Joussineau, C., Soule, J., Martin, M., Anguille, C., Montcourrier, P., and Alexandre, D. (2003). Delta-promoted filopodia mediate long-range lateral inhibition in Drosophila. Nature 426, 555–559.

Doe, C.Q., and Goodman, C.S. (1985). Early events in insect neurogenesis. II. The role of cell interactions and cell lineage in the determination of neuronal precursor cells. Dev Biol 111, 206–219.

Eldar, A., and Elowitz, M.B. (2010). Functional roles for noise in genetic circuits. Nature 467, 167–173.

Elliott, H., Fischer, R.S., Myers, K.A., Desai, R.A., Gao, L., Chen, C.S., Adelstein, R.S., Waterman, C.M., and Danuser, G. (2015). Myosin II controls cellular branching morphogenesis and migration in three dimensions by minimizing cell-surface curvature. Nat Cell Biol 17, 137–147.

Feinberg, E.H., Vanhoven, M.K., Bendesky, A., Wang, G., Fetter, R.D., Shen, K., and Bargmann, C.I. (2008). GFP Reconstitution Across Synaptic Partners (GRASP) defines cell contacts and synapses in living nervous systems. Neuron 57, 353–363.

Ferguson, J.P., Huber, S.D., Willy, N.M., Aygun, E., Goker, S., Atabey, T., and Kural, C. (2017). Mechanoregulation of clathrin-mediated endocytosis. J Cell Sci.

Fischer, R.S., Gardel, M., Ma, X., Adelstein, R.S., and Waterman, C.M. (2009). Local cortical tension by myosin II guides 3D endothelial cell branching. Curr Biol 19, 260–265.

Franke, J.D., Montague, R.A., and Kiehart, D.P. (2005). Nonmuscle myosin II generates forces that transmit tension and drive contraction in multiple tissues during dorsal closure. Curr Biol 15, 2208–2221.

Fritzsche, M., Li, D., Colin-York, H., Chang, V.T., Moeendarbary, E., Felce, J.H., Sezgin, E., Charras, G., Betzig, E., and Eggeling, C. (2017). Self-organizing actin patterns shape membrane architecture but not cell mechanics. Nat Commun 8, 14347.

Furman, D.P., and Bukharina, T.A. (2008). How Drosophila melanogaster Forms its Mechanoreceptors. Curr Genomics 9, 312–323.

Georgiou, M., and Baum, B. (2010). Polarity proteins and Rho GTPases cooperate to spatially organise epithelial actin-based protrusions. J Cell Sci 123, 1089–1098.

Gordon, W.R., Zimmerman, B., He, L., Miles, L.J., Huang, J., Tiyanont, K., McArthur, D.G., Aster, J.C., Perrimon, N., Loparo, J.J., et al. (2015). Mechanical Allostery: Evidence for a Force Requirement in the Proteolytic Activation of Notch. Dev Cell 33, 729–736.

Hadjivasiliou, Z., Hunter, G.L., and Baum, B. (2016). A new mechanism for spatial pattern formation via lateral and protrusion-mediated lateral signalling. J R Soc Interface 13.

Hamada, H., Watanabe, M., Lau, H.E., Nishida, T., Hasegawa, T., Parichy, D.M., and Kondo, S. (2014). Involvement of Delta/Notch signaling in zebrafish adult pigment stripe patterning. Development 141, 318–324.

Hartenstein, V., and Posakony, J.W. (1990). A dual function of the Notch gene in Drosophila sensillum development. Dev Biol 142, 13–30.

Heitzler, P., Bourouis, M., Ruel, L., Carteret, C., and Simpson, P. (1996). Genes of the Enhancer of split and achaete-scute complexes are required for a regulatory loop between Notch and Delta during lateral signalling in Drosophila. Development 122, 161–171.

Heusermann, W., Hean, J., Trojer, D., Steib, E., von Bueren, S., Graff-Meyer, A., Genoud, C., Martin, K., Pizzato, N., Voshol, J., et al. (2016). Exosomes surf on filopodia to enter cells at endocytic hot spots, traffic within endosomes, and are targeted to the ER. J Cell Biol 213, 173–184.

Hsu, C., Jaquet, V., Maleki, F., and Becskei, A. (2016). Contribution of Bistability and Noise to Cell Fate Transitions Determined by Feedback Opening. J Mol Biol 428, 4115–4128.

Huang, H., and Kornberg, T.B. (2015). Myoblast cytonemes mediate Wg signaling from the wing imaginal disc and Delta-Notch signaling to the air sac primordium. Elife 4, e06114.

Hunter, G.L., Hadjivasiliou, Z., Bonin, H., He, L., Perrimon, N., Charras, G., and Baum, B. (2016). Coordinated control of Notch/Delta signalling and cell cycle progression drives lateral inhibition-mediated tissue patterning. Development 143, 2305–2310.

Hutson, M.S., Tokutake, Y., Chang, M.S., Bloor, J.W., Venakides, S., Kiehart, D.P., and Edwards, G.S. (2003). Forces for morphogenesis investigated with laser microsurgery and quantitative modeling. Science 300, 145–149.

Ishizaki, T., Uehata, M., Tamechika, I., Keel, J., Nonomura, K., Maekawa, M., and Narumiya, S. (2000). Pharmacological properties of Y-27632, a specific inhibitor of rho-associated kinases. Mol Pharmacol 57, 976–983.

Karess, R.E., Chang, X.J., Edwards, K.A., Kulkarni, S., Aguilera, I., and Kiehart, D.P. (1991). The regulatory light chain of nonmuscle myosin is encoded by spaghetti-squash, a gene required for cytokinesis in Drosophila. Cell 65, 1177–1189.

Kasza, K.E., Farrell, D.L., and Zallen, J.A. (2014). Spatiotemporal control of epithelial remodeling by regulated myosin phosphorylation. Proc Natl Acad Sci U S A 111, 11732–11737.

Khait, I., Orsher, Y., Golan, O., Binshtok, U., Gordon-Bar, N., Amir-Zilberstein, L., and Sprinzak, D. (2016). Quantitative Analysis of Delta-like 1 Membrane Dynamics Elucidates the Role of Contact Geometry on Notch Signaling. Cell Rep 14, 225–233.

Kiehart, D.P., Galbraith, C.G., Edwards, K.A., Rickoll, W.L., and Montague, R.A. (2000). Multiple forces contribute to cell sheet morphogenesis for dorsal closure in Drosophila. J Cell Biol 149, 471–490.

Kiehart, D.P., Lutz, M.S., Chan, D., Ketchum, A.S., Laymon, R.A., Nguyen, B., and Goldstein, L.S. (1989). Identification of the gene for fly non-muscle myosin heavy chain: Drosophila myosin heavy chains are encoded by a gene family. EMBO J 8, 913–922.

Kornberg, T.B. (2017). Distributing signaling proteins in space and time: the province of cytonemes. Curr Opin Genet Dev 45, 22–27.

Kornberg, T.B., and Roy, S. (2014). Communicating by touch–neurons are not alone. Trends Cell Biol 24, 370–376.

Koster, D.V., Husain, K., Iljazi, E., Bhat, A., Bieling, P., Mullins, R.D., Rao, M., and Mayor, S. (2016). Actomyosin dynamics drive local membrane component organization in an in vitro active composite layer. Proc Natl Acad Sci U S A 113, E1645–1654.

Kress, H., Stelzer, E.H., Holzer, D., Buss, F., Griffiths, G., and Rohrbach, A. (2007). Filopodia act as phagocytic tentacles and pull with discrete steps and a load-dependent velocity. Proc Natl Acad Sci U S A 104, 11633–11638.

Langridge, P.D., and Struhl, G. (2017). Epsin-Dependent Ligand Endocytosis Activates Notch by Force. Cell 171, 1383–1396 e1312.

Le Gall, M., De Mattei, C., and Giniger, E. (2008). Molecular separation of two signaling pathways for the receptor, Notch. Dev Biol 313, 556–567.

Lehmann, M.J., Sherer, N.M., Marks, C.B., Pypaert, M., and Mothes, W. (2005). Actin- and myosin-driven movement of viruses along filopodia precedes their entry into cells. J Cell Biol 170, 317–325.

Leijnse, N., Oddershede, L.B., and Bendix, P.M. (2015). Helical buckling of actin inside filopodia generates traction. Proc Natl Acad Sci U S A 112, 136–141.

Liu, J., Sun, Y., Oster, G.F., and Drubin, D.G. (2010). Mechanochemical crosstalk during endocytic vesicle formation. Curr Opin Cell Biol 22, 36–43.

Losick, R., and Desplan, C. (2008). Stochasticity and cell fate. Science 320, 65–68.

Loubery, S., and Gonzalez-Gaitan, M. (2014). Monitoring notch/delta endosomal trafficking and signaling in Drosophila. Methods Enzymol 534, 301–321.

Lowe, N., Rees, J.S., Roote, J., Ryder, E., Armean, I.M., Johnson, G., Drummond, E., Spriggs, H., Drummond, J., Magbanua, J.P., et al. (2014). Analysis of the expression patterns, subcellular localisations and interaction partners of Drosophila proteins using a pigP protein trap library. Development 141, 3994–4005.

Luca, V.C., Kim, B.C., Ge, C., Kakuda, S., Wu, D., Roein-Peikar, M., Haltiwanger, R.S., Zhu, C., Ha, T., and Garcia, K.C. (2017). Notch-Jagged complex structure implicates a catch bond in tuning ligand sensitivity. Science 355, 1320–1324.

Macia, E., Ehrlich, M., Massol, R., Boucrot, E., Brunner, C., and Kirchhausen, T. (2006). Dynasore, a cell-permeable inhibitor of dynamin. Dev Cell 10, 839–850.

Major, R.J., and Irvine, K.D. (2005). Influence of Notch on dorsoventral compartmentalization and actin organization in the Drosophila wing. Development 132, 3823–3833.

Medeiros, N.A., Burnette, D.T., and Forscher, P. (2006). Myosin II functions in actin-bundle turnover in neuronal growth cones. Nat Cell Biol 8, 215–226.

Meloty-Kapella, L., Shergill, B., Kuon, J., Botvinick, E., and Weinmaster, G. (2012). Notch ligand endocytosis generates mechanical pulling force dependent on dynamin, epsins, and actin. Dev Cell 22, 1299–1312.

Mooren, O.L., Galletta, B.J., and Cooper, J.A. (2012). Roles for actin assembly in endocytosis. Annu Rev Biochem 81, 661–686.

Munjal, A., and Lecuit, T. (2014). Actomyosin networks and tissue morphogenesis. Development 141, 1789–1793.

Musse, A.A., Meloty-Kapella, L., and Weinmaster, G. (2012). Notch ligand endocytosis: mechanistic basis of signaling activity. Semin Cell Dev Biol 23, 429–436.

Palani, S., and Sarkar, C.A. (2012). Transient noise amplification and gene expression synchronization in a bistable mammalian cell-fate switch. Cell Rep 1, 215–224.

Parks, A.L., Shalaby, N.A., and Muskavitch, M.A. (2008). Notch and suppressor of Hairless regulate levels but not patterns of Delta expression in Drosophila. Genesis 46, 265–275.

Ploscariu, N., Kuczera, K., Malek, K.E., Wawrzyniuk, M., Dey, A., and Szoszkiewicz, R. (2014). Single molecule studies of force-induced S2 site exposure in the mammalian Notch negative regulatory domain. J Phys Chem B 118, 4761–4770.

Sagar, Prols, F., Wiegreffe, C., and Scaal, M. (2015). Communication between distant epithelial cells by filopodia-like protrusions during embryonic development. Development 142, 665–671.

Sayyad, W.A., Amin, L., Fabris, P., Ercolini, E., and Torre, V. (2015). The role of myosin-II in force generation of DRG filopodia and lamellipodia. Sci Rep 5, 7842.

Schweisguth, F. (2004). Regulation of notch signaling activity. Curr Biol 14, R129–138.

Shergill, B., Meloty-Kapella, L., Musse, A.A., Weinmaster, G., and Botvinick, E. (2012). Optical tweezers studies on Notch: single-molecule interaction strength is independent of ligand endocytosis. Dev Cell 22, 1313–1320.

Trylinski, M., Mazouni, K., and Schweisguth, F. (2017). Intra-lineage Fate Decisions Involve Activation of Notch Receptors Basal to the Midbody in Drosophila Sensory Organ Precursor Cells. Curr Biol 27, 2239–2247 e2233.

Tseng, Q., Duchemin-Pelletier, E., Deshiere, A., Balland, M., Guillou, H., Filhol, O., and Thery, M. (2012). Spatial organization of the extracellular matrix regulates cell-cell junction positioning. Proc Natl Acad Sci U S A 109, 1506–1511.

Vasquez, C.G., Tworoger, M., and Martin, A.C. (2014). Dynamic myosin phosphorylation regulates contractile pulses and tissue integrity during epithelial morphogenesis. J Cell Biol 206, 435–450.

Verdier, V., Guang Chao, C., and Settleman, J. (2006). Rho-kinase regulates tissue morphogenesis via non-muscle myosin and LIM-kinase during Drosophila development. BMC Dev Biol 6, 38.

Wang, W., and Struhl, G. (2004). Drosophila Epsin mediates a select endocytic pathway that DSL ligands must enter to activate Notch. Development 131, 5367–5380.

Wang, X., and Ha, T. (2013). Defining single molecular forces required to activate integrin and notch signaling. Science 340, 991–994.

Xie, X., Cho, B., and Fischer, J.A. (2012). Drosophila Epsin’s role in Notch ligand cells requires three Epsin protein functions: the lipid binding function of the ENTH domain, a single Ubiquitin interaction motif, and a subset of the C-terminal protein binding modules. Dev Biol 363, 399–412.

Zhou, R., Mohr, S., Hannon, G.J., and Perrimon, N. (2013). Inducing RNAi in Drosophila cells by transfection with dsRNA. Cold Spring Harb Protoc 2013, 461–463.

